# Gene editing of *Nicotiana benthamiana* architecture for space-efficient production of recombinant proteins in controlled environments

**DOI:** 10.1101/2025.10.01.679797

**Authors:** Béatrice Giroux, Kristina LeBreux, Louis Feyzeau, Marie-Claire Goulet, Charles Goulet, Dominique Michaud

## Abstract

Indoor vertical farming (VF) offers several practical advantages for the cultivation of plant protein bio-factories, including plant uniformity, product consistency, water/nutrient recycling and production cycles on a year-round basis. Much progress has been achieved in recent years toward the development of innovative systems for artificial lighting, automated irrigation, plant handling, environment control and space use optimization in VF systems. Here, we used a CRISPR-Cas9 gene editing approach to generate mutant lines of transient protein expression host *Nicotiana benthamiana* presenting a compact, space-efficient phenotype compared to the so-called LAB strain commonly used for protein production. Our strategy consisted of altering apical dominance by suppressing the biosynthesis of strigolactone, a negative regulator of axillary bud outgrowth-promoting cytokinins. Strigolactone-depleted lines were generated by knocking-down the expression of either *Carotenoid cleavage dioxygenase 7* (CCD7) or *Carotenoid cleavage dioxygenase 8* (CCD8), two key enzymes of the metabolic pathway leading to strigolactone synthesis. Knocking-down the genes of either enzyme had no impact on the overall growth rate of the plant but drastically influenced its leaf proteome, auxin/cytokinin ratio and overall architecture. More specifically, the ΔCCD mutants exhibited altered glycolytic and malate-processing enzyme fluxes driving the production of pyruvate and cytokinins in leaf tissue, an axillary growth-oriented development pattern and, most importantly, a spatial footprint reduced by 45% to 50% compared to the LAB strain. Most importantly, recombinant protein yields per plant were maintained in the mutant lines, as here illustrated for the model protein GFP and for rituximab, a chimeric monoclonal antibody of confirmed clinical value in humans. Our data demonstrate the usefulness of ΔCCD7 and ΔCCD8 knockout leading to strigolactone depletion for the generation of compact, space-efficient *N. benthamiana* lines well suited to VF systems intended for biopharmaceutical production.

## Introduction

Vertical farming (VF), as a complement to conventional agricultural and greenhouse systems, is rapidly emerging as an essential component of the global plant production sector, especially in urban areas (Petrovics and Giezen, 2021; Oh and Lu, 2023; Yuan et al., 2025). This new form of agriculture relies on indoor, multilayer systems in which plant growth factors such as light, temperature, humidity, atmospheric CO_2_, water and mineral nutrients are precisely controlled, independent of natural light and other outdoor conditions (SharathKumar et al., 2020). By providing an optimized environment for plant growth, VF systems offer several practical benefits, including plant uniformity, product consistency, efficient recycling of water and nutrients, production on a year-round basis, soilless cultivation with no impact on land use, and reduced pathogen or pest pressures (Gorczyca et al., 2025; Yuan et al., 2025).

Mostly seen as a sustainable solution to food security (van Delden et al., 2021), VF is also gaining interest for the production of non-food plant-derived compounds such as plant natural metabolites of pharmaceutical and nutraceutical interest, or clinically valuable biopharmaceuticals, such as vaccines and therapeutic antibodies, expressed in plant protein biofactories (Huebbers and Buyel, 2021; Stiles et al., 2025). A notable advantage of closed VF systems is the feasibility of fine-tuning growth conditions tailored to modulate metabolic fluxes and maximize the accumulation of desired compounds in the plant (Yeshi et al., 2022). Additional benefits of VF systems include their contained nature and reliance on automation, which together enable strictly controlled environments helpful in minimizing plant contamination from external sources, facilitating the use of biological agents for pest control, and easing regulatory burdens related to recombinant DNA technologies or the approval of plant-based bioproducts for clinical use (Stiles et al., 2025).

Significant progress has been made, and much remains to be done, to harness the full potential of VF systems for plant production (SharathKumar et al., 2020; van Delden et al., 2021). In recent years, multidisciplinary research efforts have enabled the engineering of systems for the fine control of lighting conditions and key environmental parameters such as temperature and air circulation in VF settings (Folta, 2019). Research progress has also contributed to the design of innovative systems for plant irrigation, the development of vertical stacking strategies for space optimization, the implementation of robotics tools for plant handling and harvesting, and the integration of artificial intelligence processes for real-time monitoring and control of growth parameters (van Delden et al., 2021; Rathor et al., 2024). As a next step to further consolidate the position of VF in the plant production sector, research must now focus on improving the physiological and morphological characteristics of the plant itself which represents, as both the integrator of growth factor signals and producer of the final product, a central determinant of the success or failure of any VF endeavor (Folta, 2019).

Currently, plants used for food production or for the manufacture of non-food products in VF facilities are selected from commercial varieties or standardized lines optimized for outdoor or greenhouse environments but that may lack traits relevant to the specific attributes of multilayer indoor settings (Teo and Yu, 2024). Over the years, crops have been bred, selected and/or optimized for yield preformance under a range of agronomic conditions and environmental constraints. By comparison, adaptability to variable environments appears less critical in controlled indoor systems, which now opens the possibility of developing ‘designer crops’ that fully exploit the favorable conditions provided by VF technologies (Folta, 2019; Stiles et al., 2025). Several traits have been identified as potentially advantageous for VF, including an enhanced photosynthetic efficiency under artificial lighting, improved light penetration in high-density plant canopies and root systems optimized for water and nutrient uptake in hydroponic setups (Teo and Yu, 2024; Stiles et al., 2025). Another trait of particular relevance is the spatial footprint, or volumetric occupancy, of the plant (Folta, 2019), which determines how many plants can be accommodated in the room and, ultimately, the yield obtained per production cycle.

In this study, we addressed the question of plant spatial footprint in the context of biopharmaceutical production, using the transient protein expression host *Nicotiana benthamiana* as a model (Akher et al., 2025). This small Solanaceae plant, as the most efficient and most versatile plant protein biofactory, has become a gold standard in both academia and industry for the high-yield production of vaccine antigens, therapeutic antibodies and other proteins of industrial interest (Eidenberger et al., 2023; Washida et al., 2025). While *N. benthamiana* production platforms often rely on plants grown in greenhouse settings, VF facilities are also used in some instances, notably in the private sector (Fukuzawa et al., 2024). Highly automated, climate-independent VF shows obvious advantages over greenhouse systems in terms of plant productivity, product uniformity and revenue predictability that could make it the preferred option of molecular farming companies in forthcoming years (Huebbers and Buyel, 2021) and trigger research efforts toward the development of *N. benthamiana* lines adapted to the specific characteristics of VF systems. CRISPR/Cas9 gene editing approaches have recently been used in recent years to improve recombinant protein yield or quality in *N. benthamiana* by the alteration of specific enzymatic or cellular functions in leaf tissue (e.g., Singh et al., 2022; Kogelmann et al., 2024, 2025; Blumberg et al., 2025; Göritzer et al., 2025). Here, we devised a CRISPR/Cas9 editing approach to generate *N. benthamiana* lines that present a compact, space-efficient phenotype compared to the original laboratory (or LAB) strain commonly used for recombinant protein expression (Bally et al., 2018).

## Results and discussion

### Rationale for the engineering of compact *N. benthamiana* lines

The aboveground architecture of higher plants is determined by the pattern of shoot branching, which itself depends on developmental cues, environmental stimuli, sugar signals and hormonal interplays (Teichmann and Muhr, 2015; Barbier et al., 2019). Despite its complexity, the regulation of shoot branching can be explained by a simple model in which three classes of hormones – auxins, cytokinins and strigolactones – interact either to promote axillary bud outgrowth or, conversely, to inhibit this process (**Figure 1A**). At the gene level, a key regulatory hub in the branching process is BRANCHED1 (or BRC1), a pivotal bud-outgrowth-inhibiting member of the TCP transcription factor family (Wang et al., 2019). Cytokinins, by repressing BRC1 expression and strigolactone biosynthesis (Chen et al., 2023), promote bud outgrowth and shoot branching, whereas strigolactones inhibit these processes through their inducing effect on BRC1 expression and their negative effects on cytokinin biosynthesis and structural integrity (Duan et al., 2019; Mashiguchi et al. 2021). Auxins do not directly affect BRC1 expression but contribute to the regulation of shoot branching through their inducing effect on the strigolactone pathway and their downregulating effect on cytokinin biosynthesis (Barbier et al., 2019; Mashiguchi et al., 2021). Along with sugar availability in axillary buds (Kebrom, 2017; Da Cao et al., 2023), these hormonal interplays between auxins, cytokinins and strigolactones shape the overall architecture of the plant. From an applied perspective, cytokinins and strigolactones, given their central roles as regulators of various plant growth processes including bud outgrowth (Li et al., 2021; Khuvung et al., 2022), are now considered as promising genetic targets for crop improvement (Nguyen et al., 2021; Mandal et al., 2022; Kelly et al., 2023).

**Figure 1.**
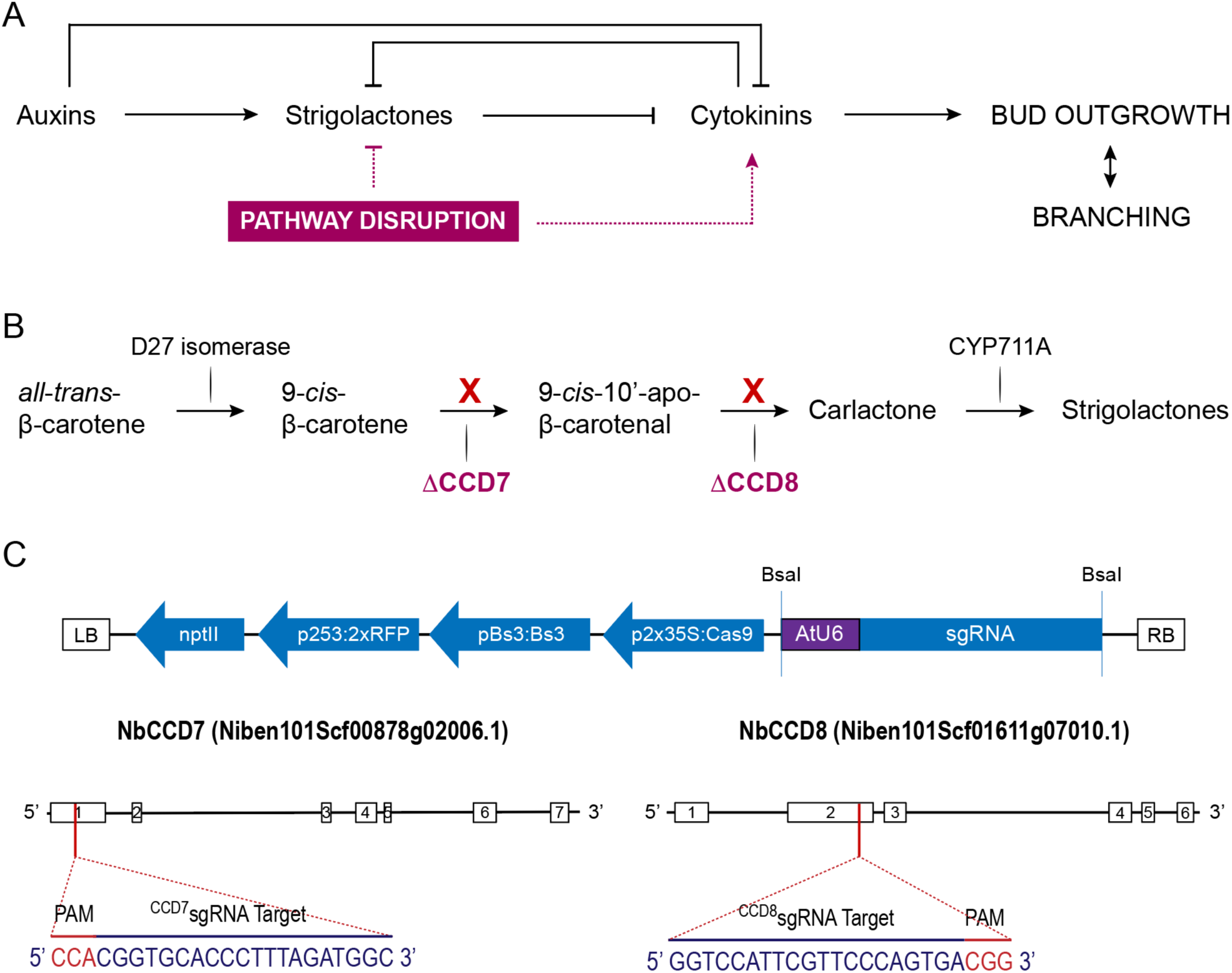
Rationale for the development of compact *N. benthamiana* lines by CRISPR/Cas9 gene editing. **A** Regulation of axillary bud outgrowth and shoot branching by auxins, strigolactones and cytokinins. Arrowheads indicate a positive interaction, flatheads a negative interaction. **B** Schematic representation of the strigolactone biosynthetic pathway. Red X marks indicate reactions targeted to produce the CCD7-depleted (ΔCCD7) or CCD8-depleted (ΔCCD8) mutant lines by CRISPR/Cas9 editing. **C** DNA constructs used to knock out the CCD7- and CCD8-encoding genes. Nucleotide sequences correspond to sgRNAs specific to target regions in the first exon of *NbCCD7* (ΔCCD7 lines) or the second exon of *NbCCD8* (ΔCCD8 lines). White boxes highlight the 6 (or 7) exons, and black lines the 5 (or 6) introns, of the two target genes. BsaI, restriction sites for sgRNA cloning; LB, left border of the T-DNA region; RB, right border of the T-DNA region; PAM, protospacer adjacent DNA motif for Cas9 binding.

Based on this, our approach to produce compact lines of *N. benthamiana* consisted of compromising strigolactone biosynthesis by knocking out key enzymes required for the production of carlactone, the immediate precursor of strigolactones along the strigolactone biosynthetic pathway (Seto et al., 2014). Strigolactones derive from the isomerization of a carotenoid precursor, *all*-*trans*-β-carotene, into 9-*cis*-β-carotene by the action of the iron-binding isomerase D27, followed by the two-step cleavage of this intermediate into carlactone by the non-heme iron enzymes CAROTENOID CLEAVAGE DIOXYGENASE 7 (CCD7) and CAROTENOID CLEAVAGE DIOXYGENASE 8 (CCD8) (Alder et al., 2012) (**Figure 1B**). Early studies with mutants of Arabidopsis, rice, pea and petunia confirmed the strict dependence of strigolactone biosynthesis on CCD7 and CCD8, as well as the strong inhibitory effect of strigolactones on bud outgrowth and shoot branching (Gomez-Roldan et al., 2008; Umehara et al., 2008; Drummond et al., 2009). Accordingly, tomato lines expressing an antisense RNA for *ccd7* transcripts (Vogel et al., 2010), tomato or potato lines expressing an RNAi sequence for *ccd8* (Kohlen et al., 2012; Pasare et al., 2013), and tomato or tobacco CRISPR/Cas9 mutants for *ccd7* or *ccd8* (Gao et al., 2018; Chen et al., 2023) all exhibited a branched, compact phenotype compared to their respective parent lines. Assuming a similar outcome for *N. benthamiana* mutants compromised in their ability to produce CCD7 or CCD8, we devised a CRISPR/Cas9 editing strategy to prevent the expression of either enzyme in the *N. benthamiana* LAB strain. Single guide (sg) RNAs targeting specific sequences in the first exon of *Nbccd7* (encoding CCD7) or the second exon of *Nbccd8* (encoding CCD8) were first designed (**Figure 1C**), and then used along with the Cas9 nuclease to produce NbCCD7-(ΔCCD7) and NbCCD8-deficient (ΔCCD8) mutants.

### A branched, compact phenotype for the ΔCCD7 and ΔCCD8 mutants

Transgenic lines expressing the antibiotic selection marker neomycin phosphotransferase II (NptII) were regenerated *in vitro* on kanamycin-containing medium following *Agrobacterium tumefaciens*-mediated transformation with the Cas9 nuclease and sgRNA coding sequences (**Figure 1**). T0 ΔCCD7 and ΔCCD8 mutant plantlets regenerated from independent calli were acclimated in a growth chamber and tested by PCR for the presence of the selection marker gene using *nptII*-specific DNA primers. A ∼500-bp *nptII* amplicon was amplified from the genomic DNA of all tested samples, confirming that the clones regenerated on kanamycin medium were all transformed by the bacterial vector. T1 plants were produced from the seeds of T0 plants by self-pollination, and their genomic DNA assessed by high resolution melting (HRM) analysis following PCR amplification with specific *ccd7* and *ccd8* DNA primers to identify homozygous mutant lines. Sanger sequencing was performed on homozygous lines showing the expected branched phenotype to identify mutants harboring distinct insertion or deletion events in either target gene. Three mutants for CCD7 (ΔCCD7-R, or Red line; ΔCCD7-Y, or Yellow line; and ΔCCD7-B, or Blue line), and three mutants for CCD8 (ΔCCD8-P, or Purple line; ΔCCD8-O, or Orange line; and ΔCCD8-G, or Green line), were selected for the production of a T2 progeny and the identification of transgene-free plants (**Figure 2A**).

**Figure 2.**
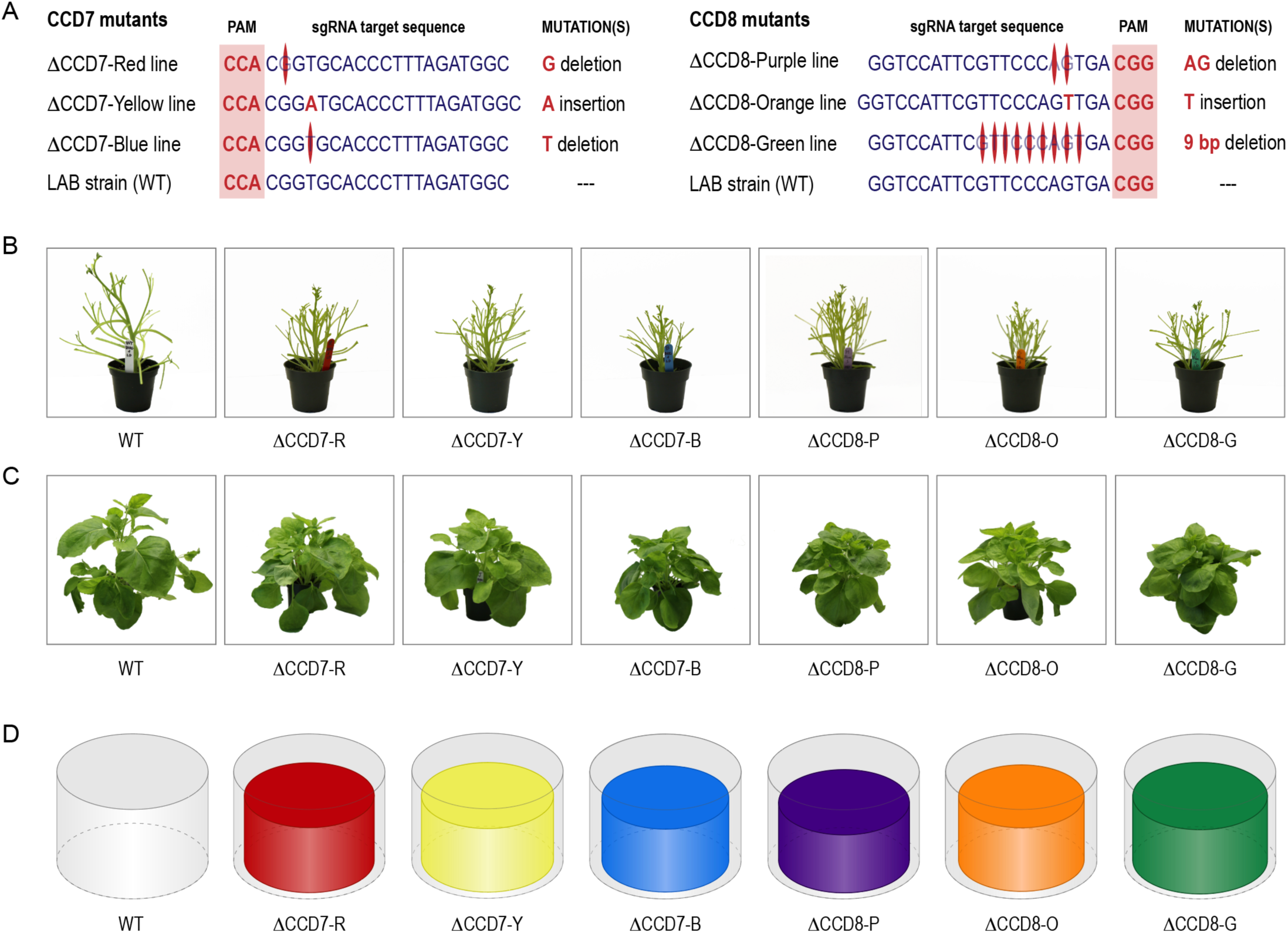
General architecture and spatial footprint of six selected Δ*Nb*CCD7 and Δ*Nb*CCD8 mutant lines. **A** CRISPR/Cas9-induced mutations in ΔCCD7-Y (Yellow), ΔCCD7-R (Red) and ΔCCD7-B (Blue) CCD7-depleted lines; and ΔCCD8-P (Purple), ΔCCD8-O (Orange) and ΔCCD8-G (Green) CCD8-depleted lines. **B** Branching patterns of the six mutants compared to the wild-type (WT) parental line. **C** Visual overview of the six mutants, compared to the WT. D. Spatial footprint (volumetric occupancy) of the six mutants (colored cylinders) compared to the WT (transparent grey cylinders). Cylinders were drawn from plant volume data presented in Table 1, relative to the WT volume (26,700 cm^3^).

**Table 1.** Growth parameters of *N. benthamiana* wild-type and strigolactone-depleted lines ΔCCD7 and ΔCCD8. ^a,b^.

| Trait | Mutant lines |  |  |  |  |  |  |
| --- | --- | --- | --- | --- | --- | --- | --- |
|  | Wild-type | ΔCCD7 |  |  | ΔCCD8 |  |  |
|  |  | ΔCCD7-Red | ΔCCD7-Yellow | ΔCCD7-Blue | ΔCCD8-Purple | ΔCCD8-Orange | ΔCCD8-Green |
| Fresh leaf biomass (g) |  |  |  |  |  |  |  |
| . Total | 45.0 ± 1.7 | 50.0 ± 1.6 | 46.6 ± 1.8 | 45.4 ± 1.6 | 50.4 ± 3.3 | 42.5 ± 2.2 | 52.4 ± 4.1 |
| . Primary stem | 33.3 ± 1.2 | 34.0 ± 0.4 | 30.6 ± 1.1 | 31.1 ± 0.8 | 33.3 ± 2.1 | 25.9 ± 1.3**<sup>c</sup> | 33.3 ± 2.1 |
| . Axillary stems | 11.8 ± 1.5 | 15.9 ± 1.3* | 16.0 ± 0.9* | 14.4 ± 1.1 | 17.1 ± 1.2** | 16.6 ± 0.9* | 18.0 ± 2.2** |
| Fresh stem biomass (g) | 24.8 ± 1.3 | 22.3 ± 0.7 | 21.4 ± 0.4 | 19.6 ± 0.5 | 21.7 ± 2.3 | 20.3 ± 1.2 | 22.3 ± 1.7 |
| Plant height (cm) | 22.8 ± 0.5 | 16.5 ± 0.3**** | 16.3 ± 0.3**** | 15.8 ± 0.3**** | 14.8 ± 0.9**** | 16.1 ± 0.2**** | 16.6 ± 0.6**** |
| Plant diameter (cm) | 38.2 ± 0.9 | 32.8 ± 0.4** | 33.3 ± 0.6** | 32.0 ± 0.9*** | 33.0 ± 1.5** | 31.7 ± 0.8*** | 34.0 ± 1.2* |
| Area per plant (cm<sup>2</sup> X 100) | 11.6 ± 0.5 | 8.5 ± 0.2*** | 8.8 ± 0.3** | 8.1 ± 0.4*** | 8.6 ± 0.8** | 7.9 ± 0.4*** | 9.1 ± 0.6** |
| Volume per plant (cm<sup>3</sup> X 1,000) | 26.7 ± 1.6 | 14.0 ± 0.4**** | 14.3 ± 0.4**** | 12.7 ± 0.8**** | 13.0 ± 1.9**** | 16.8 ± 0.7**** | 15.3 ± 1.6**** |
| Number of axillary stems | 9.5 ± 0.2 | 10.7 ± 0.4** | 10.6 ± 0.1** | 11.0 ± 0.4*** | 11.8 ± 0.1**** | 11.8 ± 0.2**** | 11.8 ± 0.1**** |
| Number of leaves |  |  |  |  |  |  |  |
| . Primary stem | 15.4 ± 0.2 | 14.9 ± 0.4 | 15.3 ± 0.3 | 15.2 ± 0.2 | 15.7 ± 0.1 | 15.8 ± 0.4 | 15.6 ± 0.3 |
| . Axillary stems | 49.4 ± 1.3 | 62.7 ± 2.7*** | 62.2 ± 0.2** | 61.6 ± 2.9** | 85.3 ± 2.1**** | 86.6 ± 1.6**** | 82.8 ± 3.6**** |
| Leaf surface per plant (cm<sup>2</sup> X 100) |  |  |  |  |  |  |  |
| . Total | 17.4 ± 0.7 | 19.0 ± 0.7 | 18.2 ± 0.5 | 17.1 ± 0.7 | 20.6 ± 1.2<sup>(P=0.065)</sup> | 18.8 ± 1.0 | 21.8 ± 1.6** |
| . Primary stem leaves | 11.1 ± 0.4 | 10.5 ± 0.3 | 9.8 ± 0.3 | 9.5 ± 0.5<sup>(P=0.063)</sup> | 10.3 ± 0.6 | 8.6 ± 0.5** | 10.7 ± 0.6 |
| . Axillary stems leaves | 6.3 ± 0.3 | 8.5 ± 0.6* | 8.4 ± 0.2* | 7.6 ± 0.5 | 10.3 ± 0.7**** | 10.2 ± 0.5**** | 11.1 ± 1.1**** |
| Floral stage at ~45 g | 5.1 ± 0.1 | 4.0 ± 0.2*** | 4.0 ± 0.3*** | 3.7 ± 0.1**** | 4.0 ± 0.2*** | 4.3 ± 0.2* | 4.0 ± 0.1*** |

<sup>a</sup> Data obtained from plants characterized 33 days after sowing. Numbers highlighted against a clear blue background were used to calculate the plant spatial footprint (volume per plant).
<sup>b</sup> Each value is the mean of four biological replicates ± SE.
<sup>c</sup> \*, $P < 0.05$ ; \*\*, $P < 0.01$ ; \*\*\*, $P < 0.001$ ; \*\*\*\*, $P < 0.0001$ (post-ANOVA Dunnett's tests with an $\alpha$ threshold for significance of 0.05).

As expected, based on the reported effects of CCD7 or CCD8 depletion in other Solanaceae species (Gao et al., 2018; Chen et al., 2023), the ΔCCD7 and ΔCCD8 mutant lines exhibited a visibly altered growth pattern suggestive of altered hormonal balances in the plant. Compared to the LAB strain, these lines displayed a branched phenotype (**Figure 2B**), with the number of axillary stems increased by almost 20%, the number of axillary stem leaves increased by 25% (ΔCCD7 lines) to 72% (ΔCCD8 lines), and the total area of axillary leaves increased by 30% (ΔCCD7 lines) to 67% (ΔCCD8 lines) (**Table 1**). The mutant lines presented a compact phenotype (**Figure 2C**), explained by a main stem height reduced by ∼30% and a ground surface area reduced by ∼27% compared to the control line (**Table 1**). Overall, these phenotypic changes led to a spatial footprint for the ΔCCD7 and ΔCCD8 lines reduced by 45-50% compared to the control line, for a similar fresh leaf biomass (∼45 g) 33 days after seed sowing (**Figure 2D**; **Table 1**).

We calculated additional growth parameters –namely the leaf harvesting index, the leaf area index, the axillary leaf area index, and the leaf/stem biomass ratio– based on the basic phenotypic data above (**Table 2**), given their potential relevance as indicators of yield or biomass quality in a plant molecular farming context (Buyel et al., 2021). The harvest index, here defined as the ratio of total leaf biomass to total plant biomass, was 7% greater for the ΔCCD7 and ΔCCD8 lines compared to the control line. The leaf area index, which refers to the total leaf area of a given plant over the area occupied by that plant, ranged from 2.08 to 2.41 for the mutant lines compared to 1.51 for the control line, indicative of a more efficient use of space per unit of cultivation area. The axillary leaf area index, here defined as the total area of axillary leaves on a given plant over the ground area occupied by that plant, was increased by 80% and 130%, respectively, for the ΔCCD7 and ΔCCD8 lines, which could possibly represent an advantage in terms of protein yield per unit of cultivation area as recombinant proteins in *N. benthamiana* accumulate preferentially in young (e.g., axillary) leaves (Goulet et al., 2019). Finally, the leaf-to-stem biomass ratio, referring to the ratio of leaf fresh biomass to stem fresh biomass, was estimated at ∼2.25 for the mutant lines compared to 1.85 for the wild-type line, a notable increase that could be advantageous upon harvest for the downstream processing of the shoot biomass (Buyel et al., 2021). Together, these additional inferences further supported the practical potential of disrupting the transcriptional integrity of the CCD7- or CCD8-encoding genes to generate *N. benthamiana* variants tailored for VF systems in terms of space use efficiency, not only at the plant canopy level but also at the individual plant level.

**Table 2.**
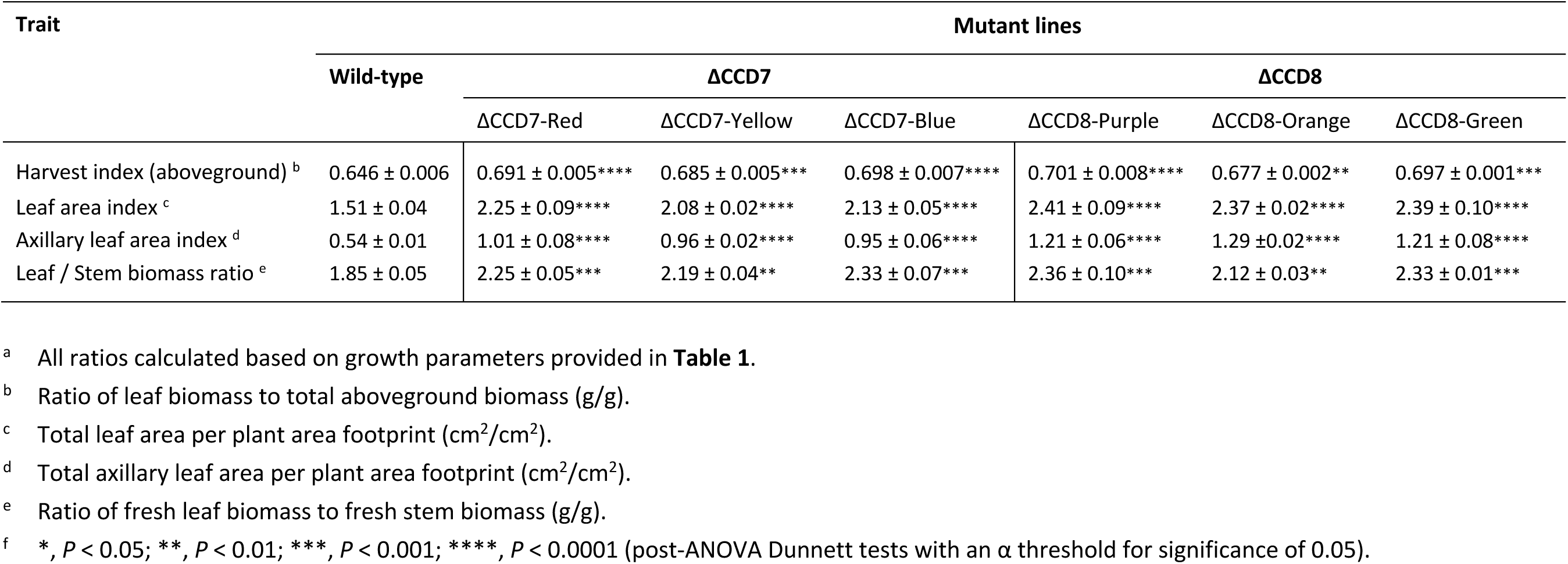
Additional growth indicators for strigolactone-depleted lines ΔCCD7 and ΔCCD8.^a^.

### An altered leaf proteome in the ΔCCD7-R and ΔCCD8-P mutants

From a physiological standpoint, a direct consequence of strigolactone depletion is an upregulation of cytokinin levels (Chen et al., 2023) and, as a result, a reduction of the auxin/cytokinin molecular ratio which influences, along with sucrose partitioning and strigolactone content, the relative strengths of apical dominance and axillary bud outgrowth (Kebrom, 2017; Da Cao et al., 2023). To confirm an alteration of the auxin/cytokinin ratio in the CCD7- and CCD8-depleted lines, we measured auxin and cytokinin contents in leaf crude extracts of line ΔCCD7–R (Red line), a ΔCCD7 mutant, and line ΔCCD8–P (Purple line), a ΔCCD8 mutant (**Figure 3**). Leaf P10, corresponding to the 10^th^ true leaf on the main stem (counting upward, from the ground), was selected for the assays based of its efficiency to produce high levels of recombinant proteins (Goulet et al., 2019; Hamel et al., 2025). As expected, given the promoting effect of cytokinins on bud outgrowth and the branched phenotype of the ΔCCD7 and ΔCCD8 mutants (see **Figure 2**), total cytokinin content was upregulated in both mutants, with relative increases of 50% in line ΔCCD7–R and 200% in line ΔCCD8–P compared to the wild-type line (**Figure 3A**). In accordance with the upstream effects of auxins on cytokinin and strigolactone biosynthesis (Barbier et al., 2019), auxin levels remained unchanged in leaf tissue, thereby resulting in reduced auxin/cytokinin ratios in the two mutants. Changes in cytokinin content were explained by increased levels of the cytokinin immediate ribosyl precursors N6-(D2-isopentenyl) adenine riboside (iPR) and *trans*-zeatin riboside (*t*ZR) (**Figure 3B**). By comparison, the mevalonate pathway-derived cytokinin *cis*-zeatin (*c*Z) and its conjugated form *cis*-zeatin riboside (*c*ZR) were barely detectable in both control and mutant lines, suggesting the upregulating effects of CCD7 and CCD8 knockdown on cytokinin content to result from an induction of the plastid-located methylerythritol 4-phosphate (MEP) pathway for iP (iPR) and *t*Z (*t*ZR) production (Kasahara et al., 2004; Sakakibara, 2025).

**Figure 3.**
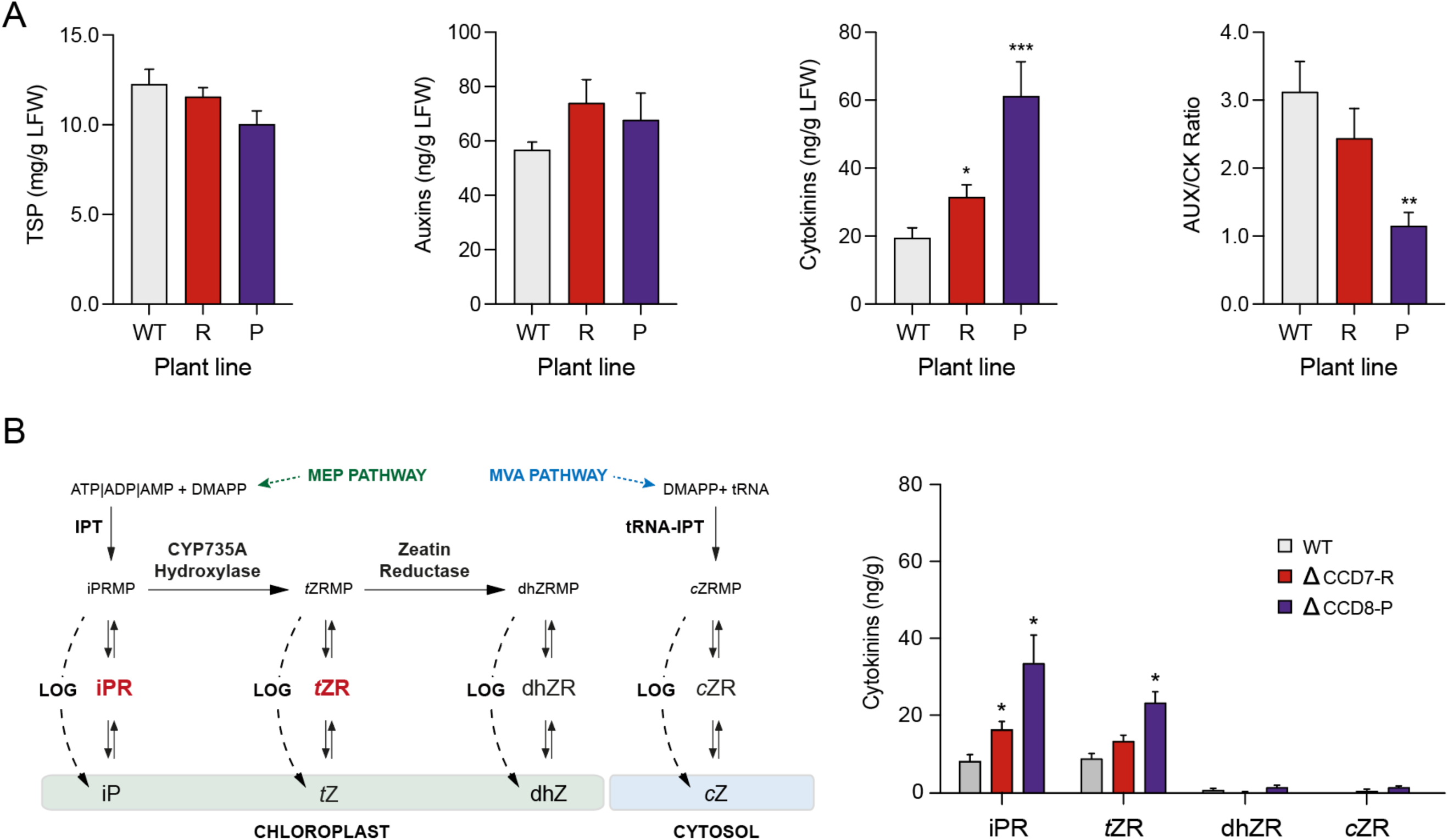
Total soluble protein, auxin and cytokinin contents in mutant lines ΔCCD7–R and ΔCCD8–P. **A** Total soluble protein, auxin and cytokinin contents in Leaf P10 of the two mutants. **B** Cytokinin biosynthetic pathways (left) and concentration of specific cytokinin forms in the two mutants (right). WT, wild-type parental line. Values on panels A and B are the mean of four biological replicates ± SE. Asterisks (*) indicate statistically significant differences compared to the WT (post-ANOVA Dunnett’s test; *p < 0.05, **p < 0.01, ***p < 0.001). DMAPP, dimethylallyl pyrophosphate; IPT, isopentenyl phosphotransferase; LOG, Lonely guy cytokinin-activating enzyme for the one-step production of cytokinin precursors (Kuroha et al., 2009); MEP pathway, methylerythritol phosphate pathway; MVA pathway, mevalonate pathway.

Assuming a wide-ranging impact of increased cytokinin levels on basic metabolic functions in leaves, we conducted an analysis of the leaf proteome to characterize the effects of CCD7 or CCD8 depletion in the ΔCCD7–R and ΔCCD8–P mutants (**Dataset S1**, available upon request; **Figure S1**). Previous studies have reported protein-inducing and protein-repressing effects for cytokinins and strigolactones at the proteome scale using plants treated with natural forms or synthetic analogs of these hormones (Zd’arska et al., 2013; Chen et al., 2014; Li et al., 2014; Berka et al., 2020; Olmedo et al., 2023). Multiple proteome alterations were also observed in mutants or transgenic lines compromised in their ability to produce active or stable forms of these same two regulators (Cerny et al., 2013; Chen et al., 2014; Li et al., 2014; Skalak et al., 2019; Pan et al., 2022), or in transgenic lines engineered to express isopentenyl transferase, an enzyme catalyzing the first, rate-limiting step of the cytokinin biosynthetic pathway (Lochmanova et al., 2010; Xu et al., 2010; Cerny et al., 2013; Skalak et al., 2016; Skalak et al., 2019; Pan et al., 2022). All in all, these studies highlighted a wide range of leaf proteome alterations in plants treated with, or modified to show altered levels of, strigolactones or cytokinins, consistent with the various roles played by these hormones at both the cellular and whole-plant levels (Li et al., 2021; Argueso and Kieber, 2024; Khuvung et al., 2022).

In accordance with the published literature, numerous alterations of the leaf proteome were detected in the ΔCCD7–R and ΔCCD8–P mutants. Out of 2,914 proteins confidently identified based on our MS/MS spectra, 770 proteins were differentially regulated in at least one mutant compared to the parent line (limma p-value < 0.05), including 352 upregulated, and 418 downregulated, proteins (**Dataset S1**, available upon request). More specifically, 219 proteins were up- or downregulated by at least 1.5-fold in one or both two mutants, including 57 upregulated proteins and 162 downregulated proteins (**Figure S1**). Of the 51 proteins upregulated by at least 1.5-fold in line ΔCCD8–P, 25 were also upregulated at this threshold in line ΔCCD7–R compared to the control line. Of the 135 proteins downregulated by at least 1.5-fold in line ΔCCD8–P, 72 were also downregulated in line ΔCCD7–R. Considering all the proteins regulated by at least 1.5-fold in at least one mutant, 31 were upregulated [and 99 downregulated] in line ΔCCD7–R, compared to 51 proteins (or 65% more) upregulated [and 135 (or 36% more) downregulated] in line ΔCCD8–P. Of the 31 upregulated proteins in line ΔCCD7–R, 25 (or 81%) were also upregulated in line ΔCCD8–P, while 72 proteins (or 73%) were downregulated in both lines, out of the 99 downregulated proteins in line ΔCCD7–R. Together, these observations confirmed the significant, wide-ranging effects of *Nbccd7* or *Nbccd8* knockdown on the leaf proteome and the onset of comparable proteome alterations in the ΔCCD7–R and ΔCCD8–P lines. Our data also suggested a dose effect of cytokinins in the two mutants, given the stronger and broader effects observed in line ΔCCD8–P which exhibited much higher contents of both the iPR and tZR cytokinin conjugates compared to the ΔCCD7–R and control lines (**Figure 3**).

### A pyruvate, energy production-oriented proteome in the ΔCCD7-R and ΔCCD8-P mutants

We performed a targeted analysis of our proteomic dataset to highlight potential effects of CCD7 or CCD8 knockdown on carbon metabolite-processing pathways in lines ΔCCD7–R and ΔCCD8–P. Proteomic studies over the last twelve years have highlighted the impact of strigolactones and cytokinins on the expression of proteins involved in primary carbon metabolism including photosynthesis, sucrose processing, glycolysis, and the tricarboxylic acid (TCA) cycle (Zd’arska et al., 2013; Chen et al., 2014; Li et al., 2014; Skalak et al., 2019; Olmedo et al., 2023). This was notably illustrated by the induction of sucrose-processing and glycolytic enzymes in plants treated with or engineered to produce more cytokinins, consistent with the roles of these hormones in controlling carbon partitioning and nutrient acquisition in sink tissues (McIntyre et al., 2021; Wu et al., 2021), and in promoting cell division (Wu et al., 2021), a highly complex process tightly linked to energy availability and fluctuations in dividing plant cells (Siqueira et al., 2018). Here, we took a close look at the relative levels of proteins involved in sucrose metabolism, glycolysis, NADH production via the TCA cycle, and oxidative phosphorylation for ATP synthesis along the mitochondrial electron transport chain (**Figure 4**; **Table S1**).

**Figure 4.**
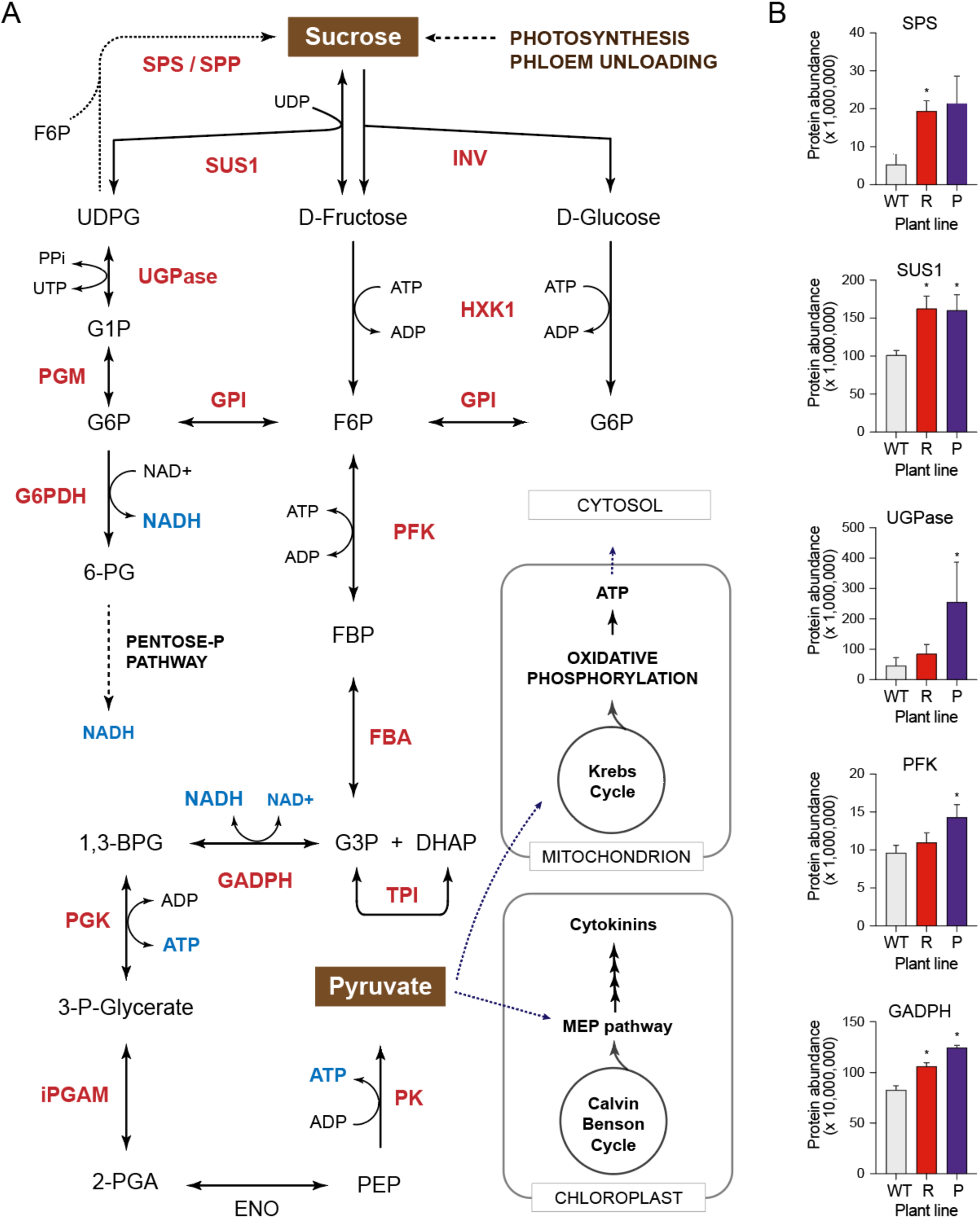
Upregulation of sucrose-metabolizing and glycolytic enzymes in Leaf P10 of mutant lines ΔCCD7–R and ΔCCD8–P. **A** Visual representation of the sucrose-processing and glycolytic pathways leading to pyruvate synthesis in the cytosol. Non-abbreviated enzyme names are provided in supplemental **Table S1**. Enzymes in red fonts were significantly upregulated in at least one mutant line, compared to the wild-type parental line (limma p-values < 0.05). **B** Relative abundance of selected sucrose-metabolizing and glycolytic enzymes upregulated in the mutant lines compared to the wild-type (WT). Values are the mean of four biological replicates ± SE. Asterisks (*) indicate statistically significant differences compared to the WT (post-ANOVA Dunnett’s test; *, p < 0.05).

In support of previous studies, most enzymes involved in sucrose metabolism and glycolysis were upregulated in at least one of the two mutants, including 13 enzymes in the cytokinin-rich ΔCCD8–P line and 4 enzymes in the ΔCCD7–R line, out of 14 enzymes identified along this segment of the primary metabolism (limma p-value < 0.05) (**Figure 4A**, **Table S1**). Some of these enzymes, such as sucrose-phosphate synthase and UDP-glucose pyrophosphorylase (**Figure 4B**), were upregulated by more than 3-fold in at least one mutant, and 8 of them were found at levels at least 1.5-fold their level in the wild-type line (**Table S1**), again suggesting comparable proteome alterations in the two mutants and a dose response associated with cytokinin content in leaf tissue. As for sucrose-processing and glycolytic enzymes in the cytosolic compartment, most mitochondrial proteins involved in the TCA cycle or oxidative phosphorylation were upregulated in the mutant lines (limma p-value < 0.05) (**Table S1**). Again, the observed effects were more pronounced in the ΔCCD8–P line, with 13 proteins significantly upregulated out of 14 assigned to the two processes, compared to 6 proteins in the ΔCCD7–R line. Consistent with the expected effects of a high cytokinin content in leaf tissue, these observations suggested a general upregulation of sugar-metabolizing, glycolytic and energy production processes in the strigolactone-depleted lines.

Our observations also highlighted the importance of pyruvate as a pivotal carbon intermediate in the CCD-depleted lines, serving not only as a carbon source to feed the TCA cycle in the mitochondrion, but also as a precursor for cytokinin production in the chloroplast via the MEP pathway (Sakakibara, 2025) (**Figure 4**). An earlier study assessing the metabolome of an Arabidopsis line engineered to overexpress a bacterial isopentenyl transferase highlighted an accelerated consumption of pyruvate in leaf tissue, accompanied by a depletion of sucrose, glucose and glucose-6-phosphate likely caused by an increased glycolytic rate to meet a higher pyruvate demand (Cerny et al., 2013). Another source of pyruvate contributing to cytokinin production in the mutant lines could involve malate, a redox carrier converted to oxaloacetate by malate dehydrogenase or to pyruvate by the malic enzyme (Dao et al., 2021). To address this possibility, we examined our proteomic data to detect potential contrasting alterations of the leaf proteome in these lines implicating malate dehydrogenase and the malic enzyme (**Figure 5**, **Table S2**). Four malate dehydrogenase isoforms and six malic enzyme isoforms could be identified in our MS/MS dataset, distributed across different cellular compartments (**Table S2**). Supporting the idea of a malate contribution to the increased demand for pyruvate, a malate dehydrogenase isoform residing in the peroxisome was strongly downregulated in the two mutant lines, and the other three isoforms, located in the cytosol, the chloroplast or the mitochondrion, were downregulated in at least one mutant (**Figure 5A**,**B**, **Table S2**). By comparison, no change was observed for the malic enzyme in line ΔCCD7–R but all six isoforms were significantly upregulated in line ΔCCD8–P. Together, these observations pointed to a general, albeit moderate, adjustment of the malate dehydrogenase/malic enzyme balance likely leading to more pyruvate in leaf cells of the ΔCCD lines, especially in line ΔCCD8–P which accumulated cytokinins at the highest levels (**Figure 3**).

**Figure 5.**
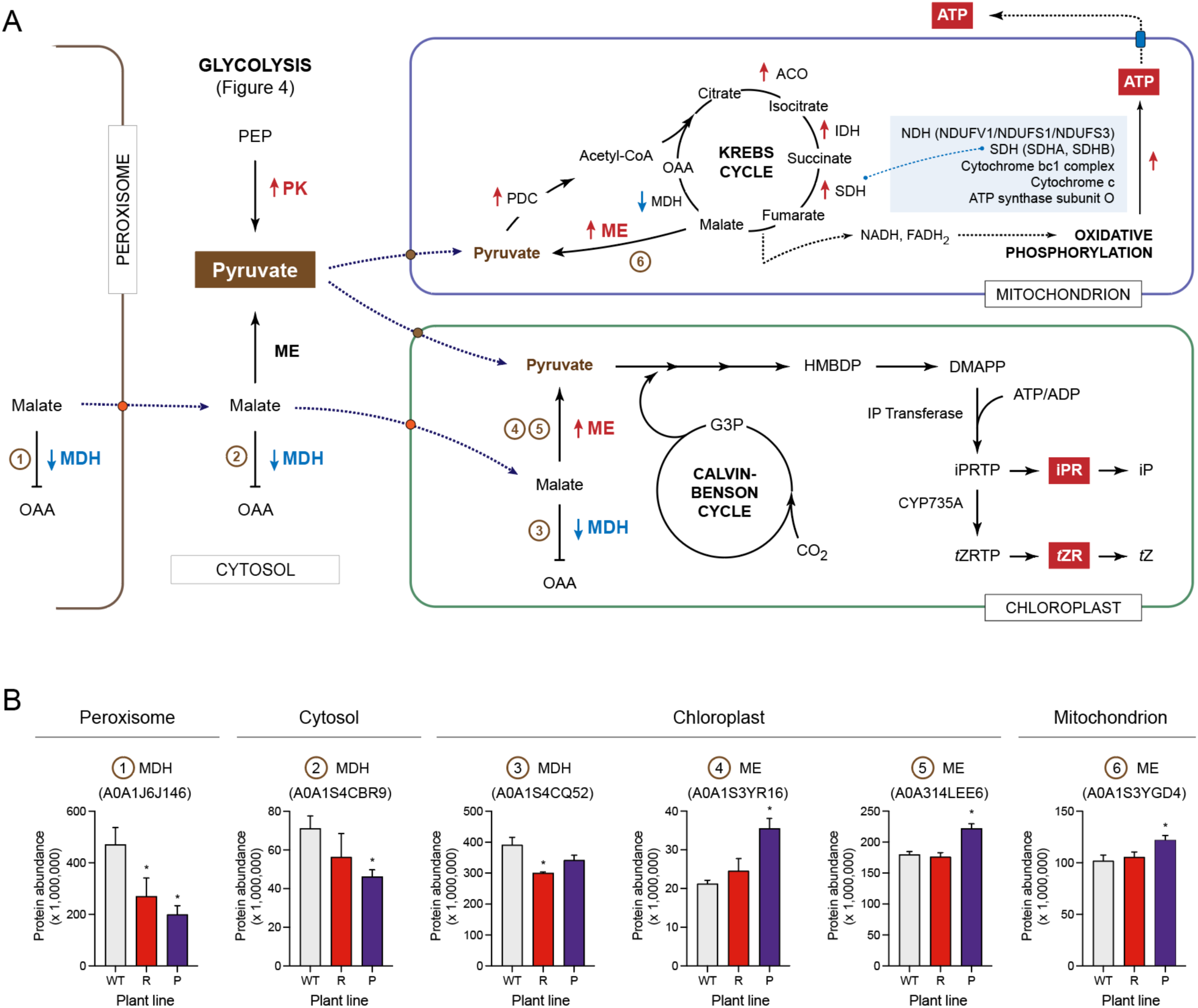
Contrasting regulation of malate dehydrogenase (MDH) and malic enzyme (ME) in Leaf P10 of mutant lines ΔCCD7–R and ΔCCD8–P. **A** Visual representation of MDH- and ME-mediated malate transformations in plant cell compartments. Arrows highlight enzyme isoforms that were significantly up- or downregulated in at least one mutant compared to the WT line (limma p-values < 0.05). Upward arrows indicate an upregulation, downward arrows a downregulation. Circled numbers next to the regulated isoforms match those used on panel B. **B** Relative abundance of MDH and ME isoforms in the two mutant lines compared to the wild-type line (WT). Each value is the mean of four biological replicates ± SE. Asterisks (*) indicate statistically significant differences compared to the control line (post-ANOVA Dunnett’s test; *, p < 0.05). More details on regulation trends for the different MDH and ME isoforms can be found in supplemental **Table S2**.

### Unaltered recombinant protein yields in the ΔCCD lines

Transient protein expression assays were conducted with the ΔCCD lines to assess their efficiency in producing recombinant proteins (**Figure 6**, **Figure 7**, **Figure S2**). Previous attempts to ectopically induce dwarf phenotypes in tobacco or *N. benthamiana* supported the potential of this approach to accommodate a larger number of plants in protein production settings (Nagatoshi et al., 2015; Rahman et al., 2024), but the variants produced showed reduced growth and leaf biomass production rates likely to compromise the net yield gains eventually obtained. By comparison, the ΔCCD mutants developed in this study not only displayed a compact architecture reducing their spatial footprint (**Figure 2**) but also maintained leaf biomass yields and growth indicators comparable to, or even better than, those of the wild-type line (**Table 1**, **Table 2**). Consistent with this, the ΔCCD7-R and ΔCCD8-P lines agroinfiltrated to express pHluorin (Jutras et al., 2018), a pH-sensitive variant of the green fluorescent protein (GFP), produced yields comparable to the wild-type, both in terms of yield per gram of fresh leaf tissue and total yield per plant (**Figure 6A**,**B**). Similarly, yields per plant and per gram of leaf tissue were similar in the mutant lines expressing rituximab (**Figure 7B**,**C**, **Figure S2B**,**C**), a chimeric monoclonal antibody used to treat autoimmune diseases and certain types of cancers (https://www.drugs.com/monograph/rituximab.html).

**Figure 6.**
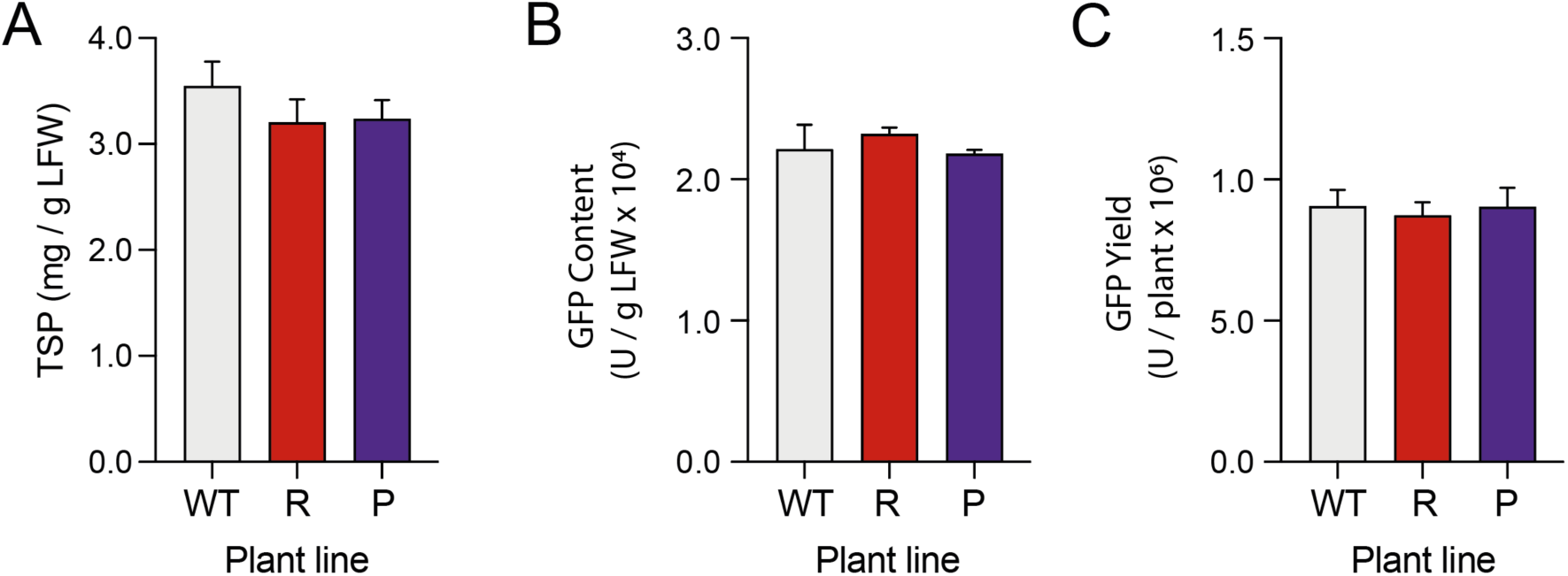
Transient expression of GFP variant pHluorin in wild-type parental line (WT) and mutant lines ΔCCD7–R and ΔCCD8–P. **A** Total soluble protein (TSP) content at the end of the expression period. **B** GFP content par gram leaf fresh weight. **C** GFP yield per plant. Samples were harvested 6 days post-infiltration. Values are the mean of four biological replicates ± SE. All mean values in this experiment were found to be statistically similar (ANOVA; p > 0.05).

**Figure 7.**
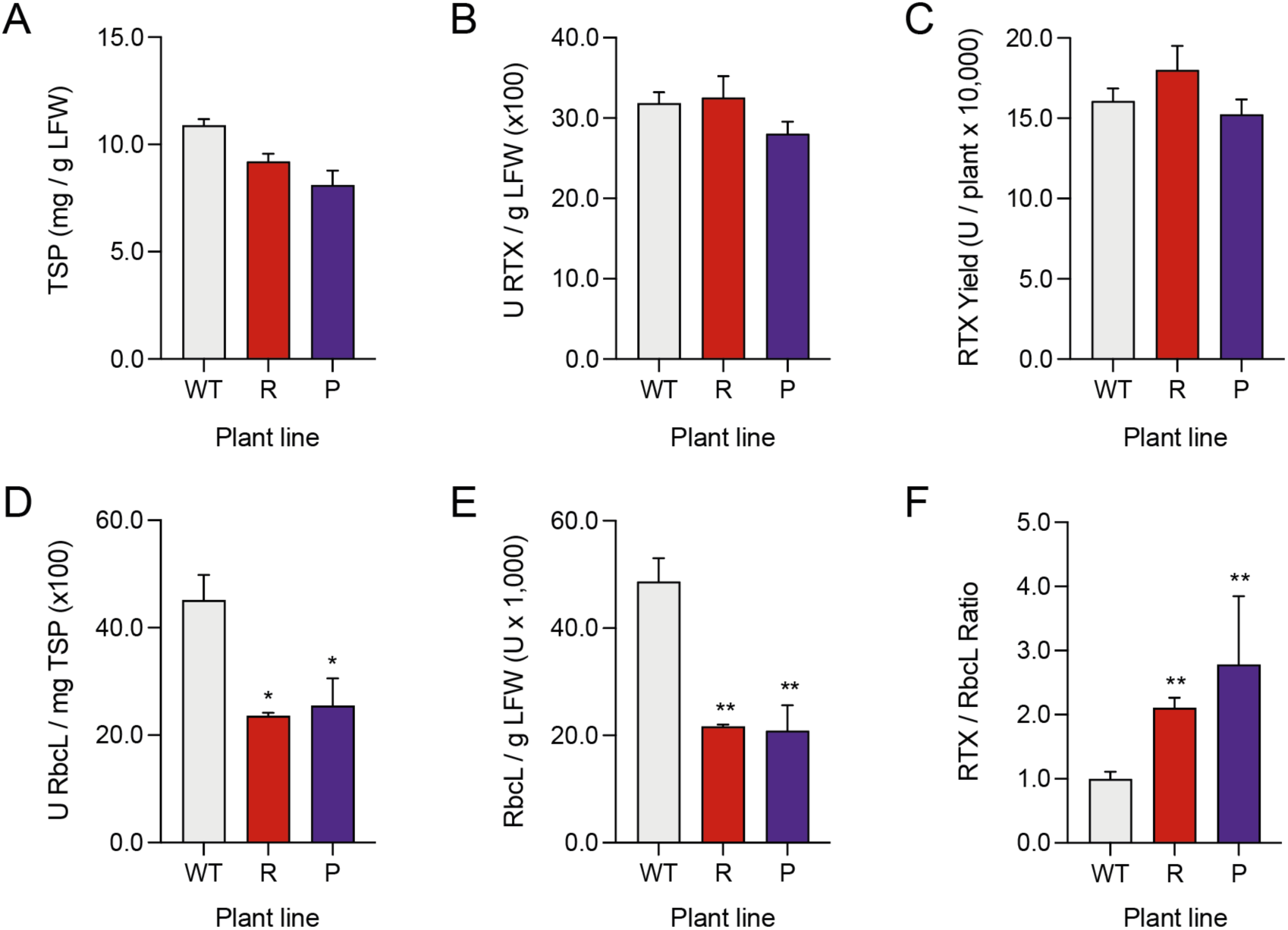
Transient expression of chimeric monoclonal antibody rituximab (RTX) in wild-type parental line (WT) and mutant lines ΔCCD7–R and ΔCCD8–P. **A** Total soluble protein (TSP) content per gram leaf fresh weight at the end of the expression period. **B** Rituximab content per gram leaf fresh weight. **C** Rituximab yield per plant. **D** Specific content of Rubisco large subunit (RbcL) per mg TSP. **E** Ponderal content of RbcL per gram leaf fresh weight. **F** Rituximab/RbcL ratio in agroinfiltrated leaves. All samples were harvested 6 days post-infiltration. Values on this figure are the mean of three or four biological replicates ± SE. Asterisks (*) indicate statistically significant differences compared to the WT (post-ANOVA Dunnett’s test; *, p < 0.05 / **, p < 0.01). Transient expression yields for all six ΔCCD7 and ΔCCD8 selected lines (Figure 2) are provided in supplemental **Figure S2**.

Unexpectedly, the specific and ponderal contents of ribulose-1,5-bisphosphate carboxylase/oxygenase (Rubisco) large subunit were reduced by more than 50% in leaves of all six ΔCCD mutants 6 days post-infiltration (**Figure 7D**,**E**, **Figure S2D**,**E**), despite roughly comparable levels of total soluble proteins at the whole-plant scale (**Figure 7A**, **Figure S2A**) or in Leaf P10 of lines ΔCCD7–R and ΔCCD8–P at the end of the growth period before infiltration (**Figure 3A**). As a result, the recombinant antibody was enriched by 100 to 150% compared to Rubisco in the ΔCCD mutant lines, prior to harvesting for downstream processing and purification (**Figure 7F**, **Figure S2F**). Rubisco, as a ‘superabundant protein’ in leaf tissue, is often regarded as a contaminant in crude extracts used for recombinant protein production, and various chromatographic or ultrafiltration approaches have been proposed to selectively remove it from crude leaf preparations following harvesting (Buyel, 2024). In this context, the intrinsic enrichment effect observed in the ΔCCD mutants could represent an additional advantage, by reducing the need for time-consuming pre-purification steps or the application of costly chemical treatments, such as the defense elicitor methyl jasmonate, to lower Rubisco levels before leaf harvesting (Robert et al., 2015).

## Conclusion

Our goal in this study was to develop strigolactone-depleted *N. benthamiana* lines showing a compact phenotype better suited to VF systems. Increasing the planting density of *N. benthamiana* canopies may already prove advantageous for improving recombinant protein yields on a per-area basis (Fujiuchi et al., 2017). On the other hand, the net gains obtained may not be optimal, given the negative impact of a high-planting density on plant growth and yield potential (Postma et al., 2020). Limited light penetration in dense canopies induces a set of adaptive responses, collectively referred to as the shade avoidance syndrome, that reduces growth and yield potential in plants shaded by their neighbors (Martinez-Garcia and Rodriguez-Concepcion, 2023). We previously reported a net yield gain of 9% per cultivation unit area for the recombinant protein hemagglutinin H1, a viral antigen used in influenza vaccines (D’Aoust et al., 2010), when expressed in agroinfiltrated *N. benthamiana* plants grown at 47 plants/m^2^ compared to a control treatment at 31 plants/m^2^ (Shang et al., 2018). This notable, but limited yield increase at the highest density, associated with a decreased leaf biomass per plant, could be partly mitigated by the use of LED inter-lighting devices (Shang et al., 2018). Given recent advances in the development of VF technologies, it is expected that lighting strategies and tools optimized for VF systems will in coming years help fill the gap between theoretical gain projections and actual yields, especially when combined with novel VF-optimized plants such as those described in this study. The ΔCCD lines, with an average area footprint reduced by 27% compared to the LAB strain (**Table 1**), could theoretically accommodate 37% more plants per production unit area while maintaining the same inter-plant spacing within the canopy. Work is ongoing to assess protein yields in the ΔCCD lines under different planting densities and lighting regimes. Work is also underway to investigate the Rubisco-depleting effect observed in these mutants upon rituximab expression, keeping in mind the recently documented impacts of agroinfiltration and defense–metabolism trade-offs on recombinant protein yields in *N. benthamiana* (Jutras et al., 2020; Kopertekh 2024; Hamel et al., 2024, 2025; Dodds et al., 2025).

## Experimental procedures

### DNA constructs for gene editing

Plasmid constructs for CRISPR/Cas9 gene editing were assembled to edit the DNA coding sequences of *N. benthamiana Carotenoid Cleavage Dioxygenase 7* (*NbCCD7*) (Niben101Scf00878g02006.1) and *N. benthamiana Carotenoid Cleavage Dioxygenase 8* (*NbCCD8*) (Niben101Scf01611g07010.1). sgRNAs were designed using the CRISPR-P v. 2.0 tool (http://crispr.hzau.edu.cn/cgi-bin/CRISPR2/CRISPR) (Liu et al., 2017), with the following target sequences: 5’–CCA CGG TGC ACC CTT TAG ATG GC–3’ (sgRNA-*NbCCD7*), located in the first exon of *NbCCD7*, or 5’–GGT CCA TTC GTT CCC AGT GAC GG–3’ (sgRNA-*NbCCD8*), located in the second exon of *NbCCD8* (**Figure 1C**) and the DNA sequence of an NbCCD8-encoding, nonfunctional pseudogene (Niben101Scf01056g05003.1). Plasmid pDGE331 harboring the Arabidopsis U6 promoter was used as shuttle vector for either sgRNAs, and plasmid pDGE463 carrying expression cassettes for Cas9, the CcdB toxin and the NptII selection marker as a recipient vector for plant transformation (AddGene) (Stuttmann et al., 2021). sgRNA cassettes were inserted by Golden Gate cloning into the recipient vector by *ccdB* excision with BsaI. The assembled vectors were cloned into *Escherichia coli* strain DH5α, extracted with the QIAprep Spin Miniprep Kit (QIAGEN), and verified by Sanger sequencing with the M13F-47 primer to confirm insertion of the sgRNA cassette. Confirmed plasmids were transferred by electroporation into *A. tumefacien*s, strain ABI, for plant transformation.

### Gene-edited ΔCCD lines

Gene-edited plant lines were generated by *Agrobacterium*-mediated transformation using 1-cm^2^ leaf explants from the *N. benthamiana* LAB strain (Bally et al., 2018). Transformed leaf cells were selected with the NptII selection marker for kanamycin resistance, and the transgenic plantlets regenerated *in vitro* as previously described (Gantner et al., 2019). Regenerated T0 plantlets were acclimated for 14 days in a PGR15 growth chamber (Conviron), under a 26°C/24°C day–night temperature cycle, a 16:8 h light/dark photoperiod, and a light intensity of 200 µmol m^-2^ s^-1^. Relative humidity in the growth chamber was maintained for two weeks at 80% using a lid placed over the plantlets, and then progressively reduced at 50% following the acclimation period. Integration of the *NptII* transgene in kanamycin-resistant plants was confirmed by PCR using genomic DNA extracted from acclimated plants, with the following primers for amplification: 5’– ACTGAAGCGGGAAGGGACTGGCTGCTATTG and 3’–GATACCGTAAAGCACGAG-GAAGCGGTCAG, producing a ∼500-base DNA amplicon after electrophoresis in 1% (w/v) agarose gels. T1 plants were obtained by manual self-pollination of NptII-confirmed T0 plants, and their genomic DNA analyzed by high resolution melting (HRM) after PCR amplification to identify homozygous mutants. The following primers were used for amplification: *ΔCCD7*-HRM-F (5’-AGAAGCCACTGAACCCATCA-3’)/*ΔCCD7*-HRM-R (5’-TAACCTGACCTGAACCACCA-3’) for *ΔCCD7*; and *ΔCCD8*-HRM-F (5’-GTTCTAATAAAGTGAGGGGACGTA-3’)/*ΔCCD*8-HRM-R (5’-GGAAGAAGGAACAAATGAGAGG-3’) for *ΔCCD8*. Sanger sequencing was performed on homozygous lines presenting a branched phenotype to identify mutants harboring distinct insertion/deletion (indel) events. The following primer sets were used for sequencing: *ΔCCD7*-SEQ-F (5’-CCATGTACCACGAGCCATAA-3’)/*ΔCCD7*-SEQ-R (5’-GGGCTTCAAGATTTGAGCAG-3’) for *ΔCCD7*; and *ΔCCD8*-SEQ-F (5’-CGCGTCCCTAACTGATAACG-3’)/*ΔCCD*8-SEQ-R (5’-GTGCAATAGTACCTTGCATCG-3’) for Δ*NbCCD8*. Six mutant lines –three ΔCCD7 and three ΔCCD8 lines– displaying an increased shoot branching phenotype with no apparent delay in leaf biomass production were selected to produce T2 plants and isolate transgene-free clones.

### Assessment of growth parameters

Basic growth parameters were monitored under greenhouse conditions to assess growth patterns and biomass production performance of the ΔCCD7 and ΔCCD8 mutants compared to the wild-type parental line. Experiments were conducted at the High-Performance Greenhouse Complex of Université Laval, Québec City QC, Canada (46°46ʹ N, 71°16ʹ W). Gene-edited and wild-type seeds were soaked in water for 30 h, sown in seedling trays, and placed for 3 days under a dome in a PGR15 growth chamber (Conviron) at 28°C, with a 16 h light/8 h dark photoperiod, 200 µmol m^-2^ s^-1^ light intensity and 75% relative humidity. The dome was removed after 3 days, and the seedlings grown for an additional 11 days under the same conditions. During the first week, seedlings were subirrigated with water for 5-min periods; in the second week, they received a 1.0 mS/cm Plant-Prod 12-2-14 Optimum Water Soluble Fertilizer supplemented with the Plant-Prod Chelated Micronutrient Fertilizer Mix (Plant Products, Laval QC, Canada) under the same irrigation regime. Young plants were transferred to the greenhouse on day 15 and grown for approximately 3 weeks at a density of 36 plants/m^2^, under a 26°C/24°C day-night temperature cycle, continuous light (24 h light/0 h dark), 160 µmol m^-2^ s^-1^ light intensity, and 60% relative humidity. The plants were irrigated using drip tubes, with a 1.6 mS/cm fertilizing solution during the first week, and with a 2.6 mS/cm fertilizing solution until leaf infiltration 33 to 35 days after sowing, once the plants had reached 40-45 g fresh leaf biomass. On the day of agroinfiltration, three plants from each mutant or control line were randomly selected to assess growth parameters, including plant height and diameter, primary and axillary leaf biomass, the number of primary and axillary stems and leaves, and the total area of primary and axillary leaves.

### Protein extraction, SDS-PAGE and immunoblotting

Leaf samples for protein, proteome and hormonal measurements were disrupted with 2.8-mm zirconium ceramic oxide beads in a Bead Ruptor 24 homogenizer (Omni International). Soluble proteins were extracted in two volumes (w/v) of ice-cold 50 mM Tris-HCl buffer, pH 8.0, containing 500 mM NaCl, 1 mM phenylmethylsulfonyl fluoride (Biobasic) and 2 mM sodium metabisulfite (Sigma-Aldrich). The homogenates were centrifuged at 4°C for 10 min at 20,000 × g. Total soluble proteins in the supernatant were quantified using the Bradford protein assay (Bio-Rad), with bovine serum albumin as a standard (Sigma-Aldrich). Electrophoretic separations by SDS-PAGE or non-reducing SDS-PAGE were performed at 200 V using a Bio-Rad Mini PROTEAN 3**®** electrophoresis system (Bio-Rad). Proteins were transferred on Hybond ECL nitrocellulose membranes (GE Healthcare) using a MiniTrans-Blot Electrophoretic Transfer Cell**®** (Bio-Rad), according to the manufacturer’s instructions.

### Leaf proteome analyses

Leaf proteomes were analyzed by ultra-high-performance liquid chromatography-tandem mass spectrometry (UHPLC-MS/MS) at the Plateforme protéomique du Centre hospitalier universitaire de Québec (Québec, Canada). Protein extracts from Leaf P10 samples of four biological replicates per line (wild-type [LAB] line, line ΔCCD7–R and line ΔCCD8–P) were prepared as follows: proteins were precipitated overnight at -20°C with five volumes of cold acetone, centrifuged, and resuspended by sonication in 50 mM ammonium bicarbonate containing 1% (w/v) sodium deoxycholate. Resuspended proteins (20 µg) were reduced with dithiothreitol, alkylated with iodoacetamide, and digested with Sequencing Grade Trypsin (Promega) at a protease/protein ratio of 1:50. Digestion was stopped by acidification with 50% (v/v) formic acid, and the resulting peptides were desalted on C18 StageTips prior to LC-MS/MS analysis. The samples were loaded on a Thermo Vanquish Neo UHPLC system (Thermo Fisher Scientific) equipped with an Evosep capillary column (EV1109, 8 cm × 150 µm ID, 1.5 µm particle size) heated to 45°C. Peptides were separated over 21 min along a linear gradient of 2% to 30% (v/v) acetonitrile containing 0.1% (v/v) formic acid, at a flow rate of 1 µl/min. The column was connected to an EASY-Spray™ source equipped with a 30-µm stainless steel emitter interfaced with an Orbitrap Exploris™ 480 MS (Thermo Fisher Scientific). Spray voltage was set at 1,900 V, and the heated capillary maintained at 280°C. Full MS spectra were acquired in profile mode, and the DIA scans collected in centroid mode under positive polarity. An internal lock-mass calibration (EASY-IC) was applied at the beginning of each run. Full MS resolution was set to 60,000, with an AGC target of 300% and a maximal IT set to 25 ms. The mass peptide range was set at 300–1500, and the DIA performed with 27 isolation windows of 20 *m/z* covering the 350-890 *m/z* precursor range, at a resolution of 15,000 with an AGC target of 800% and a maximal IT of 40 ms. The normalized HCD collision energy was set at 30%.

Raw data were processed in DIA-NN v. 1.9.2 to extract DIA signals and quantify peptides. A prediction library generated from a *Nicotiana* FASTA file (TaxID 4085) was used, with the following search criteria: a maximum of two trypsin missed cleavages, carbamidomethylated cysteines (fixed modifications), oxidized methionines (variable modifications) and N-terminal methionine excision (variable modifications). Proteins were identified based on a False Discovery Rate threshold of 1% and quantified using the MaxLFQ algorithm implemented in the diann R package. Missing values were imputed with the 1st percentile intensity (per sample). Only proteins with ≥75% of intensity values in at least one group and supported by ≥2 peptides were retained for quantification.

### Hormone quantifications

Auxin and cytokinin concentrations in crude protein extracts of Leaf P10 (see above) from four biological repetitions were quantified by High-Performance Liquid Chromatography-Electrospray Ionization (HPLC-ESI)-MS/MS at the Plant Hormone Profiling Facility, National Research Council of Canada, Aquatic and Crop Resource Development Research Center (Saskatoon SK, Canada) as previously described (Lulsdorf et al., 2013). Analyses were performed using the Multiple Reaction Monitoring function of the MassLynx v. 4.1 software (Waters). Chromatographic traces were processed off-line in QuanLynx v. 4.1 (Waters), wherein each peak was integrated and the resulting ratio of signals (nondeuterated/internal standard) compared with a calibration curve, in ng per sample. Calibration curves were generated from standard solutions based on the ratio of each analyte peak area to that of its corresponding internal standard. Quality control samples, internal standard blanks and solvent blanks were included in each batch of tissue samples.

### Binary vectors for protein transient expression assays

Protein transient expression assays were conducted to assess the efficiency of the ΔCCD lines to produce recombinant proteins, using pHluorin and rituximab as protein models. The gene construct for pHluorin (GenBank accession AF058694) was described previously (Jutras et al., 2015). The gene construct for rituximab contained expression cassettes for the light and heavy chains of the chimeric murine/human antibody (DrugBank accession no. DB00073), each fused to an N-terminal signal peptide to direct secretion in the apoplast. DNA templates were synthesized for both chains (Integrated DNA Technologies). The heavy chain coding sequence, including the signal peptide, corresponded to a plant codon-optimized version (Vector Builder Codon Optimization Tool [https://en.vectorbuilder.com]) of nucleotide Sequence 4 from Patent WO02096948 (GenBank Accession no. AX709548). The gene sequence for the light chain, also codon-optimized and including the N-terminal signal peptide, corresponded to a modified version of nucleotide Sequence 6 from the same patent (GenBank Accession no. AX709550) presenting a deletion of cytosine 525 to restore the open reading frame. The two chain coding sequences were each placed downstream of a duplicated Cauliflower mosaic virus 35 promoter, flanked with a 5’ Tobacco etch virus enhancer sequence and a 3’ nopaline synthase terminator, as described previously (Robert et al., 2013). A DNA cassette for the tumbusvirus silencing suppressor P19 (Silhavy et al., 2002) was also included, with the same promoter and regulatory elements. The resulting expression cassette was transferred into the binary vector pCambia2300 (CAMBIA) and introduced into *Agrobacterium* by electroporation for transient protein expression assays.

### Agroinfiltration and transient protein expression

Binary vectors for pHluorin and rituximab were maintained in *A. tumefaciens* strain AGL1, in the presence of appropriate antibiotics. Prior to leaf infiltration, the agrobacteria were grown to stationary phase at 28°C in Luria-Bertani medium supplemented with appropriate antibiotics, until reaching an OD_600_ of ∼4.8. The bacteria were pelleted and resuspended in infiltration buffer (10 mM MES, pH 5.6, 10 mM MgCl_2_) to an OD_600_ of 0.6. Four plants of each line (control or mutant) were vacuum-infiltrated with bacterial suspensions carrying either the pHluorin or rituximab expression cassettes. After infiltration, the plants were incubated for 7 days in a PGW40 growth chamber (Conviron) to allow for recombinant heterologous expression, under an ambient temperature of 20°C, 60% relative humidity, a 16 h day/8 h night photoperiod, a light intensity of 200 μmol m^−2^ s^−1^ and a cultivation density of 56.4 plants.m^−2^, and tap water provided as needed. Leaf samples were collected 7 days post-infiltration to quantify pHluorin and rituximab contents, relative to the wild-type parental line. GFP was quantified in 1/20 dilutions of the protein extracts using a BioTek Synergy H1 fluorimeter (Agilent Technologies), under excitation and emission wavelengths of 485 nm and 520 nm, respectively. Rituximab was detected as a ∼150-kDa band following 10% (w/w) SDS-PAGE under non-reducing conditions and quantified by in-gel densitometry with the Phoretix 2D Expression software (Nonlinear Dynamics). The large subunit of Rubisco (RbcL) was quantified by densitometry on nitrocellulose membranes using the same image analysis software, following electrophoretic separation by 12% (w/v) SDS-PAGE under reducing conditions, electrotransfer to nitrocellulose membranes and immunodetection with anti-RbcL polyclonal IgG (Agrisera) and goat anti-rabbit IgG conjugated to alkaline phosphatase (Bio-Rad). Protein–antibody complexes were visualized using the alkaline phosphatase substrate 5-bromo-4-chloro-3-indolyl phosphate and nitro blue tetrazolium for color development (Life Technologies).

### Statistical analyses

The experimental design involved 60 plants per line distributed in four groups (blocks) corresponding to different zones (or microenvironments) in the greenhouse, for a total of 4 repetitions per tested line and 420 plants overall (4 blocks × 15 plants/block × 7 lines, including the control parental line). Within each block, 3 plants were randomly selected and treated as subsamples (pseudo-replicates) for growth measurements, or pooled prior to hormone quantification and leaf proteome analyses. Of the remaining 12 plants, 3 per block were randomly selected for agroinfiltration, and their leaves pooled after leaf harvest for recombinant protein and Rubisco quantifications. Analyses of variance (ANOVA) were performed using the R software, v. 4.3.1 (R Project for Statistical Computing; https://www.r-project.org) to test for line (treatment) effects on growth, hormone and protein expression variables. Treatment means were compared using Dunnett’s tests when the ANOVA indicated a significant overall effect. An α value of 5% (0.05) was used as threshold for statistical significance.

## Acknowledgements

We thank Dr. Marc-André D’Aoust (Aramis Biotechnologies, Québec, Canada) for helpful comments on a previous version of this manuscript. This work was supported by a Discovery grant from the Natural Science and Engineering Research Council of Canada.

## Conflicts of interest

The authors declare no conflict of interest.

## Abbreviations

ANOVA: analysis of variance
CCD7: Carotenoid Cleavage Dioxygenase 7
CCD8: Carotenoid Cleavage Dioxygenase 8
MEP pathway: methylerythritol 4-phosphate pathway
TCA cycle: tricarboxylic acid cycle (Krebs cycle)
VF: vertical farming

## Funding

Natural Science and Engineering Research Council of Canada.

## Notes

### Competing Interest Statement

The authors have declared no competing interest.

