## Supplemental Figures for "Gene editing of *Nicotiana benthamiana* architecture for space-efficient production of recombinant proteins in controlled environments"

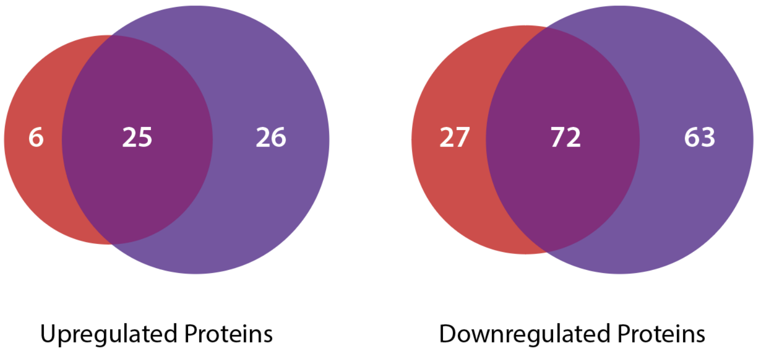


**Figure S1** Venn diagram for the numbers of proteins up- or downregulated by at least 50% in Leaf P10 of mutant lines ΔCCD7–R and ΔCCD8. Proteins considered were up- or downregulated by at least 50% compared to the wild-type (WT), as inferred from limma p-values with an α threshod of 0.05. Specific details on the regulation trends of each protein are given in supplemental **Dataset S1**, available upon request.

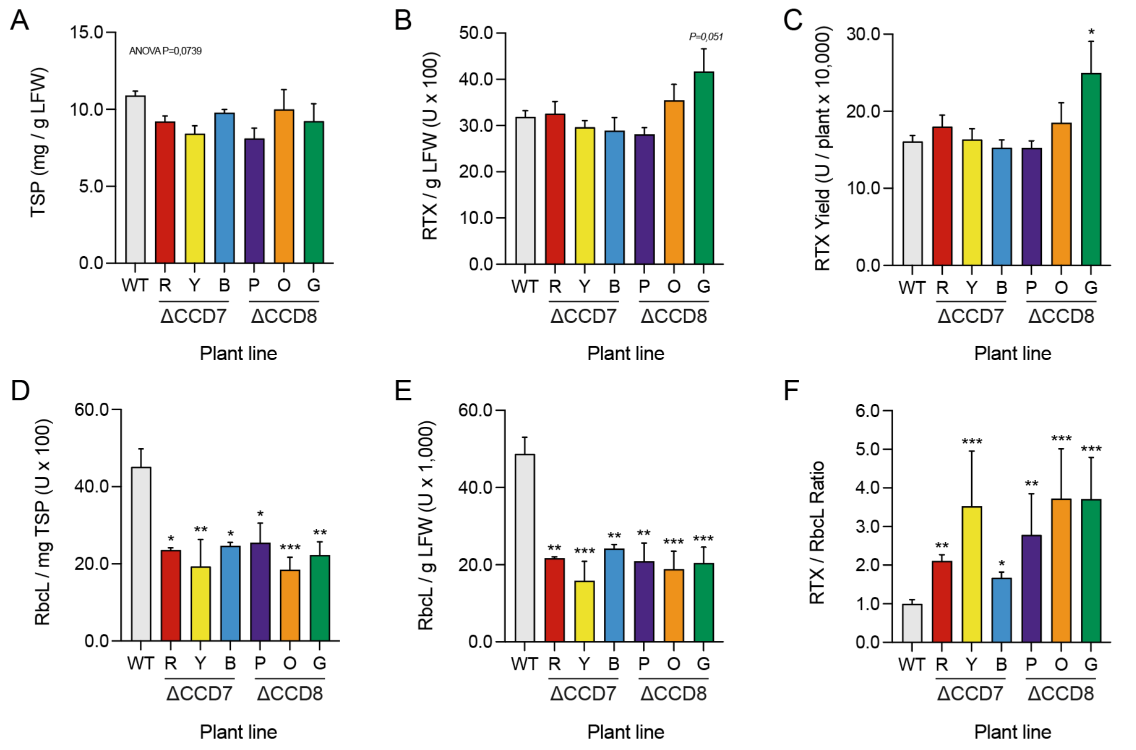


**Figure S2** Transient expression of chimeric monoclonal antibody rituximab (RTX) in wild-type parental line (WT) and mutant lines ΔCCD7–R, ΔCCD7–Y, ΔCCD7–B, ΔCCD8–P, ΔCCD8–O and ΔCCD8–G. **A** Total soluble protein (TSP) content per gram leaf fresh weight at the end of the expression period. **B** Rituximab content per gram leaf fresh weight. **C** Rituximab yield per plant. **D** Specific content of Rubisco large subunit (RbcL) per mg TSP. **E** Ponderal content of RbcL per gram leaf fresh weight. **F** Rituximab-to-RbcL ratio in agroinfiltrated leaves. All samples were harvested 6 days post-infiltration. Values on this figure are the mean of three or four biological replicates ± SE. Asterisks (*) indicate statistically significant differences compared to the WT (post-ANOVA Dunnett's test; *, $p < 0.05$ / **, $p < 0.01$ / ***, $p < 0.001$).
