## Supplemental Tables for "Gene editing of *Nicotiana benthamiana* architecture for space-efficient production of recombinant proteins in controlled environments"

**Table S1** Sugar metabolizing, glycolytic, TCA cycle and oxidative phosphorylation enzymes and proteins up- or downregulated in the ΔCCD7-R and ΔCCD8-P mutant lines.[1]

| UNIPROT No.[2] | Protein | Abbreviation | EC No. | Cell location | ΔCCD7-R[3] | ΔCCD8-P |
|---|---|---|---|---|---|---|
| | **Sucrose processing** | | | | | |
| A0A1J6I8F8 | Sucrose phosphate synthase | SPS | 2.4.1.14 | Cytosol | **1.873** | 2.015 |
| A0A1J6IAM7 | Sucrose synthase (SUS1) | SUS1 | 2.4.1.13 | Cytosol | **0.682** | **0.662** |
| A0A1S3YUU6 | UDP-glucose pyrophosphorylase | UGPase | 2.7.7.9 | Cytosol | 0.880 | **2.458** |
| A0A314LE77 | Phosphoglucomutase | PGM | 5.4.2.2 | Cytosol | 0.128 | **0.458** |
| F2Z9R3 | Glucose-6-P dehydrogenase | G6PDH | 1.1.1.49 | Cytosol | **0.426** | **0.554** |
| A0A1J6L3E7 | Invertase (β-fructofuranosidase) | INV | 3.2.1.26 | Vacuole | – | **0.321** |
| | **Glycolysis – Phase 1** | | | | | |
| Q9SEK2 | Hexokinase-1 | HXK1 | 2.7.1.1 | Cytosol | -0.312 | **0.645** |
| A0A1S4CI25 | Phosphohexose isomerase | GPI | 5.3.1.9 | Cytosol | 0.180 | **0.415** |
| A0A1S3YDQ2 | 6-phosphofructokinase (ATP-dependent) | PFK | 2.7.1.11 | Cytosol | 0.192 | **0.574** |
| A0A1J6IGW4 | Fructose-biphosphate aldolase (Aldolase) | FBA | 4.1.2.13 | Cytosol | 0.244 | **0.571** |
| | **Glycolysis – Phase 2** | | | | | |
| B3F8I0 | Glyceraldehyde-3-P dehydrogenase | GAPDH | 1.2.1.12 | Cytosol | **0.357** | **0.589** |
| A0A1S4BTZ7 | Phosphoglycerate kinase | PGK | 2.7.2.3 | Cytosol | 0.204 | **0.472** |
| A0A1S3XWD3 | Phosphoglycerate mutase | iPGAM | 5.4.2.12 | Cytosol | 0.151 | **0.282** |
| A0A1J6ITL2 | Pyruvate kinase | PK | 2.7.1.40 | Cytosol | – | **0.335** |

**Table S1** Cont'd.[1]

| UNIPROT No.[2] | Protein | Abbreviation | EC No. | Cellular location | ΔCCD7-R | ΔCCD8-P |
|---|---|---|---|---|---|---|
| | **Tricarboxylic cycle** | | | | | |
| | PYRUVATE DEHYDROGENASE COMPLEX | PDC | | | | |
| A0A1S4B2J2 | *Pyruvate dehydrogenase E1 component, subunit $\beta$* | PDHB | 1.2.4.1 | Mitochondrion | **0.214** | **0.363** |
| A0A1S4AK33 | *Pyruvate dehydrogenase E1 component, subunit $\alpha$* | PDHA | 1.2.4.1 | Mitochondrion | 0.111 | **0.222** |
| A0A1S4C175 | *Dihydrolipoamide acetyltransferase E2 component* | E2 | 2.3.1.12 | Mitochondrion | **0.498** | 0.572 |
| A0A1S3Y542 | | | | | **0.273** | **0.364** |
| | | | | | **0.260** | **0.343** |
| | | | | | **0.218** | 0.148 |
| A0A1S3WZQ8 | Aconitase hydratase (aconitase) | ACO | 4.2.1.3 | Mitochondrion | **0.259** | **0.469** |
| A0A314KP37 | | | | | **0.208** | **0.371** |
| A0A1U7W1S4 | Isocitrate dehydrogenase | IDH | 1.1.1.42 | Mitochondrion | – | **0.248** |
| A0A1J6IKJ3 | | | | | 0.156 | **0.248** |
| A0A1U7WU40 | | | | | 0.148 | **0.230** |
| A0A1S3YKI8 | Succinate dehydrogenase[4] | SDH | 1.3.5.1 | Mitochondrion | 0.082 | **0.424** |
| A0A1J6J5W3 | | | | | 0.080 | **0.416** |
| A0A1J6IUY6 | Malate dehydrogenase | MDH | 1.1.1.37 | Mitochondrion | **-0.290** | 0.097 |
| | **Oxidative phosphorylation** | | | | | |
| | **Complex I** – NADH DEHYDROGENASE | NDH | 7.1.1.2 | Mitochondrion | | |
| A0A1S4AAX1 | *Flavoprotein 1* | NDUFV1 | | | **0.448** | **0.727** |
| A0A1S4BIG4 | *Iron-sulfur protein 1* | NDUFS1 | | | **0.285** | **0.513** |
| Q5M9T3 | *Iron-sulfur protein 3* | NDUFS3 | | | 0.075 | **0.441** |

**Table S1** Cont'd.[1]

| UNIPROT No. [2] | Protein | Abbreviation | EC No. | Cell location | ΔCCD7-R | ΔCCD8-P |
|---|---|---|---|---|---|---|
| | **Oxidative phosphorylation (Cont'd)** | | | | | |
| | **Complex II** – SUCCINATE DEHYDROGENASE [4] | SDH | 1.3.5.1 | Mitochondrion | | |
| A0A1S3YKI8 | *Flavoprotein subunit* | SDHA | | | 0.082 | **0.424** |
| A0A1J6J5W3 | *Iron-sulfur subunit* | SDHB | | | 0.080 | **0.416** |
| A0A1J6J759 | **Complex III** – Cytochrome bc1 complex, sub. 7 | QCR7 (UQCRB) | N/A | Mitochondrion | 0.172 | **0.358** |
| A0A314KZN8 | Cytochrome c | Cyt c | N/A | Mitochondrion | **0.431** | **0.617** |
| A0A1S3WZA4 | **Complex V** – ATP synthase subunit O | OSCP | N/A | Mitochondrion | – | **0.339** |

[1] Data in the last two columns are expressed as log2 ratio values for either mutant compared to the control strain. Positive values indicate an upregulation in the mutant line(s), negative values a downregulation. All enzymes in the table were detected in the two ΔCCD lines and significantly regulated (**in bold**) in at least one line (limma p-values < 0.05).

[2] UniProt accession numbers retrieved from the UniProt consortium database (https://www.uniprot.org), Dec. 2024 to Mar. 2025.

[3] Absolute log2 values smaller than 0.070 (<|5%| change) were not included in the table (– sign).

[4] Enzyme located in the mitochondrial inner membrane; participates in both oxidative phosphorylation (Complex II) and the tricarboxylic cycle.

**Table S2** Malate-processing enzymes up- or downregulated in the ΔCCD7–R and ΔCCD8–P mutants.[1]

| UNIPROT No. [2] | EC Number | Cellular location [3] | ΔCCD7-R [4] | ΔCCD8-P |
|---|---|---|---|---|
| **MALATE DEHYDROGENASE** | | | | |
| A0A1J6J146 | 1.1.1.37 | Peroxisome | **-0.801** | **-1.235** |
| A0A1S4CBR9 | | Cytosol | -0.337 | **-0.625** |
| A0A1S4CQ52 | | Chloroplast | **-0.380** | -0.194 |
| A0A1J6IUY6 | | Mitochondrion | **-0.290** | 0.097 |
| **MALIC ENZYME** | | | | |
| A0A1S3YR16 | 1.1.1.40 | Chloroplast | 0.212 | **0.742** |
| A0A1S3Y729 | | | – | **0.292** |
| A0A314LEE6 | | | – | **0.305** |
| A0A1S4B7M0 | 1.1.1.39 | Mitochondrion | – | **0.242** |
| A0A1S3YGD4 | | | – | **0.258** |
| A0A1J6I8D1 | | | – | **0.250** |

1 Data in the last two columns are expressed as log2 ratio values for either mutant compared to the control strain. Positive values indicate an upregulation in the mutant line(s), negative values a downregulation. All enzyme isoforms in the table were detected in the two ΔCCD lines and significantly regulated (**in bold**) in at least one line (limma p-values < 0.05).

2 UniProt accession numbers retrieved from the UniProt consortium database (https://www.uniprot.org), Dec. 2024 to Mar. 2025.

3 As determined from available information in Uniprot for each isoform or related isoforms in Arabidopsis, and reconfirmed by sequence analysis inferences using the subcellular prediction program Deep Loc 2.0 (https://services.healthtech.dtu.dk/services/DeepLoc-2.0/).

4 Absolute log2 values smaller than 0.070 (<|5%| change) were not included the table (– sign).
